# TWO CLASSES OF MYOSIN INHIBITORS, BLEBBISTATIN AND MAVACAMTEN, STABILIZE β-CARDIAC MYOSIN IN DIFFERENT STRUCTURAL AND FUNCTIONAL STATES

**DOI:** 10.1101/2020.12.19.423544

**Authors:** Sampath K. Gollapudi, Weikang Ma, Srinivas Chakravarthy, Ariana C. Combs, Na Sa, Stephen Langer, Thomas C. Irving, Suman Nag

**Affiliations:** Department of Biochemistry, MyoKardia Inc., a wholly-owned subsidiary of Bristol Myers Squibb (TM), Brisbane, CA 94005; BioCAT, Department of Biological Sciences, Illinois Institute of Technology, Chicago, IL, USA; Department of Molecular, Cellular and Developmental Biology, BioFrontiers Institute, University of Colorado, Boulder, CO 80309, USA

**Keywords:** Super-Relaxed State (SRX), Interacting Heads Motif (IHM), Small Angle-X-ray Scattering (SAXS), mavacamten, blebbistatin, human β-cardiac myosin, Synthetic thick filaments (STF)

## Abstract

In addition to a conventional relaxed state, a fraction of myosins in the cardiac muscle exists in a newly-discovered low-energy consuming super-relaxed (SRX) state, which is kept as a reserve pool that may be engaged under sustained increased cardiac demand. The conventional and the super-relaxed states are widely assumed to correspond respectively to a structure where myosin heads are in an *open* configuration, free to interact with actin, and a *closed* configuration, inhibiting binding to actin. Disruption of the SRX population in different heart diseases, such as hypertrophic cardiomyopathy, results in unwarranted muscle contraction, and stabilizing them using myosin inhibitors is budding as an attractive therapeutic strategy. Here we examine the structure-function relationships of two myosin ATPase inhibitors, mavacamten, and blebbistatin, and found that binding of mavacamten to myosin at a site different than blebbistatin populates myosin into the SRX state. Blebbistatin, and para-nitroblebbistatin, binding to a distal pocket to the myosin lever arm near the nucleotide-binding site, does not affect the usual myosin SRX state but instead appears to render myosin into a new, perhaps non-functional, ‘ultra-relaxed’ state. X-ray scattering-based rigid body modeling shows that both mavacamten and para-nitroblebbistatin induce novel conformations that diverge significantly from the hypothetical *open* and *closed* states and furthermore, mavacamten treatment causes a greater compaction than para-nitroblebbistatin. Taken together, we conclude that mavacamten and blebbistatin stabilize myosin in different structural states, and such states may give rise to different functional energy-sparing SRX states.

## INTRODUCTION

Myosin thick filaments constitute one of the central components in the sarcomeres of striated muscles. Under activating conditions, binding of Ca^2+^ to troponin tightly regulates myosin interaction with actin on the thin filaments, which leads to sarcomere contraction (reviewed in (1, 2)). However, the discovery that in relaxed muscle, myosin can exist in two distinct structural states—a folded-back *closed* state termed as the interacting heads-motif (IHM) (3–9) and a more conventional *open* state (reviewed in (10–15))—emphasizes another mode of regulation within thick filaments that determines the number of myosin heads available for interaction with actin. In the folded back *closed* state, myosin heads are closely associated with the thick filament shaft, where one of the heads of a single myosin dimer, known as the ‘blocked head,’ has its actin-binding domain sequestered into the folded molecule, while the other head is ‘free’ (reviewed in (10–15)). Based on many different low-resolution electron microscopy structures of thick filaments and soluble myosins, it has been hypothesized that structurally, the ‘blocked head’ interacts with the converter domain of the ‘free head’ and the myosin subfragment-2 to give rise to the IHM state (reviewed in (10–15)). On the other hand, myosins in the conventional *open* state, which are thought to be in equilibrium with the *closed* state, project out from the thick filament and are ready to interact with actin under activating conditions of the muscle. Recently, in two independent studies using cryo-electron microscopy, a 4-8 Å resolution IHM structure of smooth muscle myosin has been determined (16, 17), but this is yet to become available for cardiac myosin.

As these fascinating structural discoveries were being made, parallel investigations discovered that myosin heads in relaxed thick filaments exist in an equilibrium between two biochemically-defined functional states (reviewed in (18–20)): (1) the disordered relaxed state (DRX), in which myosin has an average ATP turnover time of <60 s; and (2) the super-relaxed (SRX) state, in which myosin has a prolonged ATP turnover time of >100 s. In conjunction with the structural results, these biochemical discoveries led to a hypothesis, within the central protein structure-function dogma, that the structurally *closed* and *open* states of myosin are responsible for the low-energy consuming SRX and the high-energy consuming DRX states, respectively.

Since these discoveries, altering the distribution of myosin population between *closed* and *open* states using a small-molecule approach has emerged as an attractive therapeutic strategy to correct the aberrant cardiac function in certain cardiomyopathies. A newly discovered small molecule cardiac myosin inhibitor, mavacamten (21), which recently concluded phase-III clinical trials to treat obstructive hypertrophic cardiomyopathy, is effective in reducing the strength of muscle contraction. The ability of mavacamten to counter enhanced contractility has also been investigated *in vitro* using hypertrophic cardiomyopathy-causing mutations in myosin, myosin-binding protein C, troponin T, and troponin I (21–24). Many elegant mechanistic studies showed that, functionally, mavacamten populates myosin in the low-energy consuming SRX state (23, 25, 26). Using electron microscopy and muscle fiber X-ray diffraction studies, it has also been shown that mavacamten structurally stabilizes a *closed* state reminiscent of the IHM state (23), thereby suggesting a link between the structurally *closed* state and the SRX functionality. Whether this structure-function relationship holds, in general, is an unanswered question.

To address this question and to interrogate the mechanism of mavacamten’s action on cardiac myosin, we compared it with members of a different family of myosin inhibitors, blebbistatin, and its derivative para-nitroblebbistatin. Like mavacamten, the blebbistatin family of molecules binds and inhibits phosphate release from myosin, leading to inhibition of myosin ATPase function (27–29). In the presence of ADP, blebbistatin is also known to prime the myosin lever arm to the pre-stroke orientation (30). Structural studies using low-resolution electron microscopy and X-ray diffraction in skeletal systems (31–35) and fluorescent polarization spectroscopy in the cardiac system (36) have shown that, like mavacamten and blebbistatin can promote a folded-back myosin structure mimicking a *closed* or IHM-like state. However, single ATP turnover kinetics assessed using a single high concentration of the compound showed that, unlike mavacamten, blebbistatin does not populate more myosin in the SRX state (37). Instead, it slows the ATP turnover rates of myosins already in the SRX state (37). Altogether, these pieces of evidence suggest that although blebbistatin and mavacamten-treated myosins may be enzymatically inhibited, the mechanism of inhibition may have different functional outcomes, thereby challenging the link between the *closed* and the SRX state. Moreover, in the absence of any high-resolution atomic structures, it remains unresolved whether the *closed* structural state of myosin stabilized by blebbistatin differs from mavacamten and whether this could lead to distinct SRX functional outcomes.

In this study, using various biochemical techniques, we report that mavacamten and blebbistatin binding to different pockets of myosin lead to distinct functional outcomes. Importantly, we show that these functional differences arise from different structural states that are preferentially stabilized by mavacamten and blebbistatin. We present arguments suggesting that these two small-molecule myosin inhibitors act via different mechanisms distinguishable by dissimilar structural and functional transitions.

## RESULTS

### Mavacamten and Blebbistatin Family of Molecules Effectively Inhibit Myosin ATPase Activity

We first evaluated the concentration-dependent effects of mavacamten, blebbistatin, and para-nitroblebbistatin on basal, actin-activated, as well as myofibrillar ATPase activity of myosin (Fig. 1). Consistent with their inhibitory action, all three molecules showed potent inhibition of the basal ATPase activity of myosin in bovine cardiac synthetic thick filaments (BcSTF) (Fig. 1A), with mavacamten being the most potent among the three (IC_50_ of 0.63±0.05 μM) followed by blebbistatin (1.12±0.07 μM) and para-nitroblebbistatin (2.37±0.12 μM). A similar pattern of inhibition was also observed in the actin-activated (Fig. 1B and Table 1) and bovine cardiac myofibrillar (BcMF) ATPase activity at pCa 6 (Fig. 1C and Table 1). IC_50_ values reported for mavacamten, and blebbistatin in this study are consistent with those reported in the literature (21, 25, 26, 28, 29, 38).

**Table 1.**
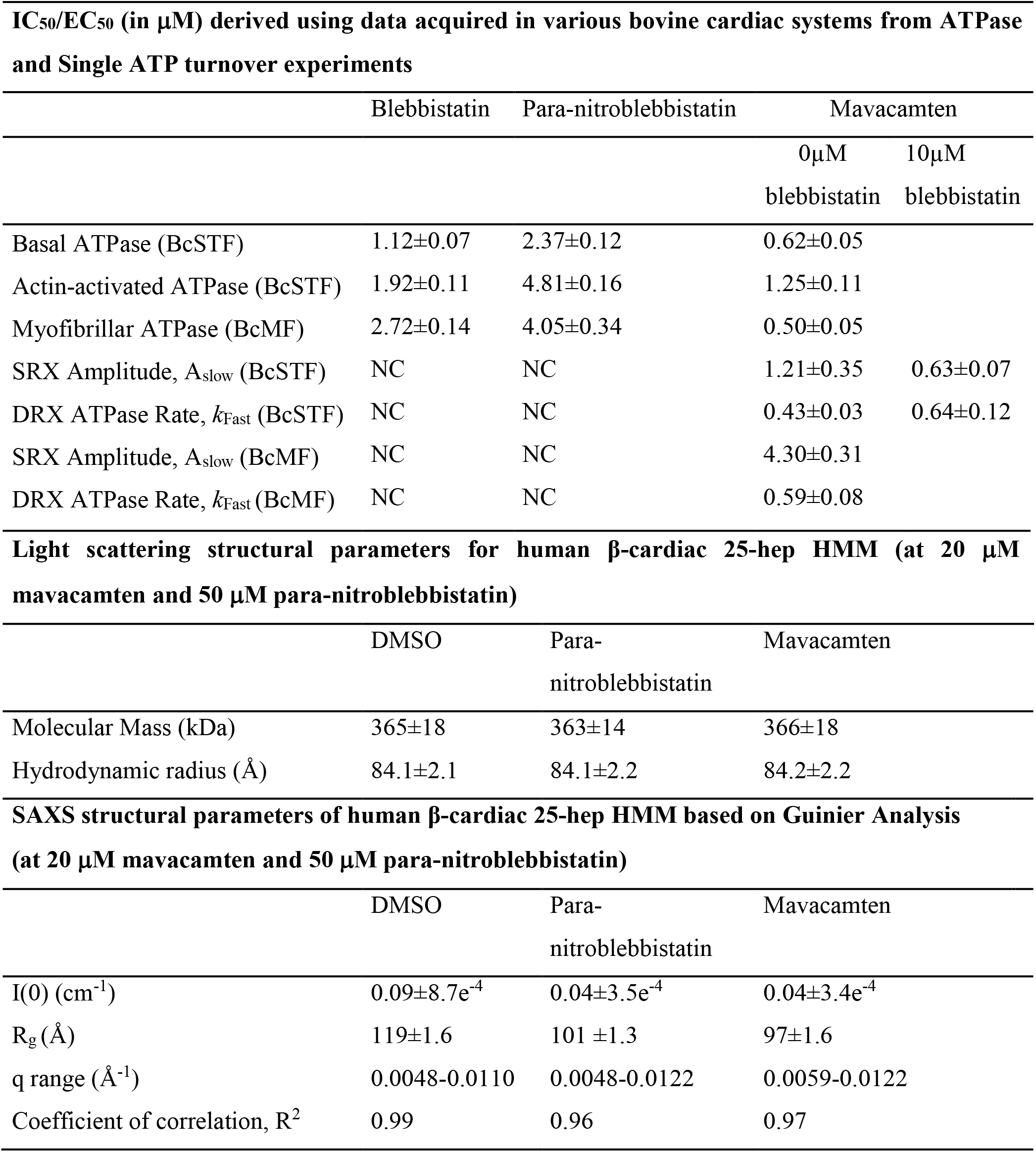
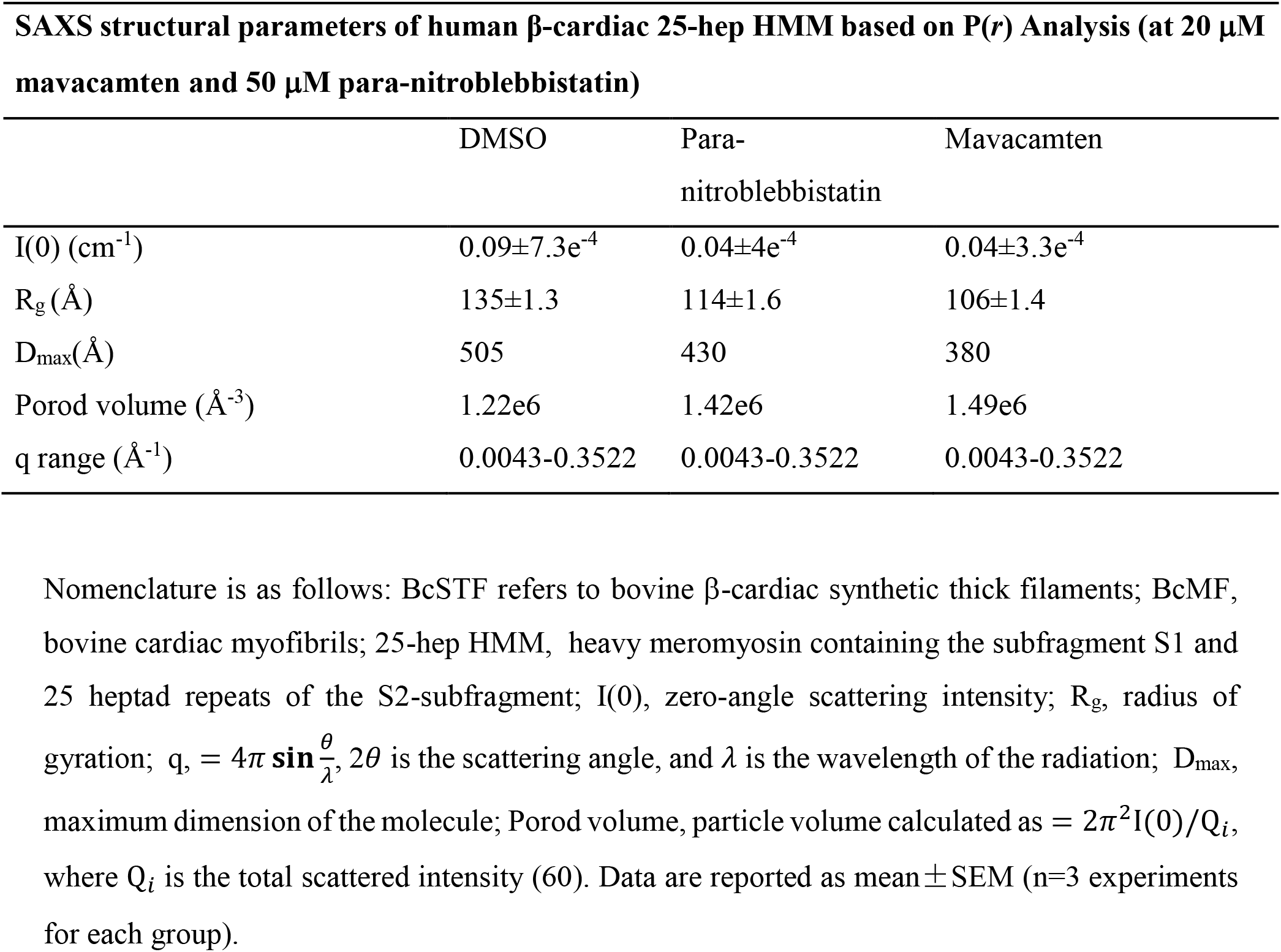
Parameters derived from ATPase, single ATP turnover and SAXS studies (NC, no change)

| IC<sub>50</sub>/EC<sub>50</sub> (in μM) derived using data acquired in various bovine cardiac systems from ATPase and Single ATP turnover experiments |  |  |  |  |
| --- | --- | --- | --- | --- |
|  | Blebbistatin | Para-nitroblebbistatin | Mavacamten |  |
|  |  |  | 0μM<br>blebbistatin | 10μM<br>blebbistatin |
| Basal ATPase (BcSTF) | 1.12±0.07 | 2.37±0.12 | 0.62±0.05 |  |
| Actin-activated ATPase (BcSTF) | 1.92±0.11 | 4.81±0.16 | 1.25±0.11 |  |
| Myofibrillar ATPase (BcMF) | 2.72±0.14 | 4.05±0.34 | 0.50±0.05 |  |
| SRX Amplitude, A<sub>slow</sub> (BcSTF) | NC | NC | 1.21±0.35 | 0.63±0.07 |
| DRX ATPase Rate, k<sub>Fast</sub> (BcSTF) | NC | NC | 0.43±0.03 | 0.64±0.12 |
| SRX Amplitude, A<sub>slow</sub> (BcMF) | NC | NC | 4.30±0.31 |  |
| DRX ATPase Rate, k<sub>Fast</sub> (BcMF) | NC | NC | 0.59±0.08 |  |
| Light scattering structural parameters for human β-cardiac 25-hep HMM (at 20 μM mavacamten and 50 μM para-nitroblebbistatin) |  |  |  |  |
|  | DMSO | Para-nitroblebbistatin | Mavacamten |  |
| Molecular Mass (kDa) | 365±18 | 363±14 | 366±18 |  |
| Hydrodynamic radius (Å) | 84.1±2.1 | 84.1±2.2 | 84.2±2.2 |  |
| SAXS structural parameters of human β-cardiac 25-hep HMM based on Guinier Analysis (at 20 μM mavacamten and 50 μM para-nitroblebbistatin) |  |  |  |  |
|  | DMSO | Para-nitroblebbistatin | Mavacamten |  |
| I(0) (cm<sup>-1</sup>) | 0.09±8.7e<sup>-4</sup> | 0.04±3.5e<sup>-4</sup> | 0.04±3.4e<sup>-4</sup> |  |
| R<sub>g</sub> (Å) | 119±1.6 | 101 ±1.3 | 97±1.6 |  |
| q range (Å<sup>-1</sup>) | 0.0048-0.0110 | 0.0048-0.0122 | 0.0059-0.0122 |  |
| Coefficient of correlation, R<sup>2</sup> | 0.99 | 0.96 | 0.97 |  |

|  | DMSO | Para-nitroblebbistatin | Mavacamten |
| --- | --- | --- | --- |
| I(0) (cm<sup>-1</sup>) | 0.09±7.3e<sup>-4</sup> | 0.04±4e<sup>-4</sup> | 0.04±3.3e<sup>-4</sup> |
| R<sub>g</sub> (Å) | 135±1.3 | 114±1.6 | 106±1.4 |
| D<sub>max</sub>(Å) | 505 | 430 | 380 |
| Porod volume (Å<sup>-3</sup>) | 1.22e6 | 1.42e6 | 1.49e6 |
| q range (Å<sup>-1</sup>) | 0.0043-0.3522 | 0.0043-0.3522 | 0.0043-0.3522 |

Nomenclature is as follows: BcSTF refers to bovine β-cardiac synthetic thick filaments; BcMF, bovine cardiac myofibrils; 25-hep HMM, heavy meromyosin containing the subfragment S1 and 25 heptad repeats of the S2-subfragment; I(0), zero-angle scattering intensity; R<sub>g</sub>, radius of gyration; $q = 4\pi \sin \frac{\theta}{\lambda}$ , $2\theta$ is the scattering angle, and $\lambda$ is the wavelength of the radiation; D<sub>max</sub>, maximum dimension of the molecule; Porod volume, particle volume calculated as $= 2\pi^2 I(0)/Q_i$ , where $Q_i$ is the total scattered intensity (60). Data are reported as mean ± SEM (n=3 experiments for each group).

**Figure 1.**
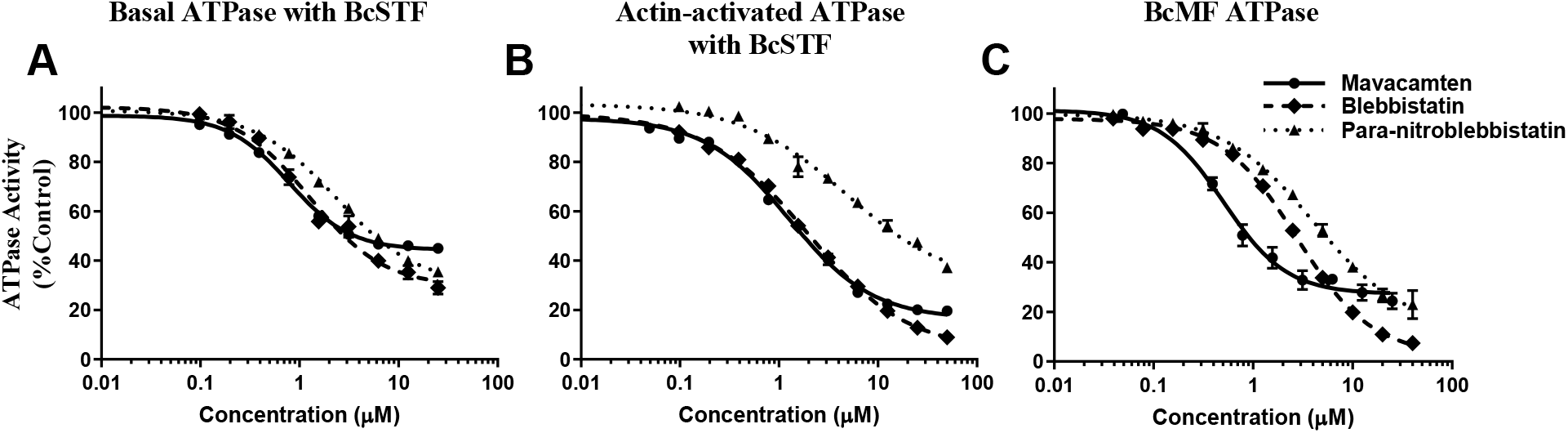
Basal, Actin-Activated, and Myofibrillar Myosin ATPase activity is Inhibited by Mavacamten, Blebbistatin, and Para-nitroblebbistatin. Normalized (**A**) Basal, (**B**) Actin-activated, and (**C**) Myofibrillar ATPase activity. The DMSO values were used to normalize the data in the respective treated systems. BcSTF refers to bovine cardiac synthetic thick filaments, and BcMF refers to bovine cardiac myofibrils. Circles, diamonds, and triangles correspond to mavacamten, blebbistatin, and para-nitroblebbistatin, respectively. The half-maximal change (IC_50_) of mavacamten estimated from these curves are shown in Table 1. Data are expressed as mean±SEM (*n*=8 from two experiments).

### Binding of Mavacamten to a Different Site on Myosin than of Blebbistatin, Populates Myosin in the SRX State

To understand if the ATPase inhibition of mavacamten, blebbistatin, and para-nitroblebbistatin arises from increasing the myosin population in the SRX state, we next studied the single ATP myosin turnover kinetics in reconstituted BcSTF in response to increasing concentrations of these three molecules. Representative mant-ATP fluorescence decay profiles after chasing with non-fluorescent ATP in these experiments are shown in Fig. 2. A qualitative comparison showed that the addition of mavacamten (Fig. 2A) led to a concentration-dependent rightward shift in the fluorescence decay profile, suggesting a slower release of mant-nucleotides from myosin. In contrast, blebbistatin (Fig. 2B) or para-nitroblebbistatin (Fig. 2C) did not show any noticeable change in the fluorescence decay up to a time point of ~ 200 sec. However, unlike mavacamten, for both blebbistatin and para-nitroblebbistatin, the fluorescence decay did not change much from 200 sec onwards (compare grey shaded region in Fig. 2A with Fig. 2B–C), and the baseline fluorescence increased in a concentration-dependent manner (grey shaded region Fig. 2B–C). A comparison of the total area under these fluorescence decay curves shows a concentration-dependent increase in the rate of undissociated mant-nucleotides from myosin in the presence of mavacamten but not blebbistatin and para-nitroblebbistatin (Fig. 2D). However, the baseline fluorescence increase for both blebbistatin and para-nitroblebbistatin treatment suggests a small but significant portion of myosins are bound to unreleased nucleotide products even after 2 hrs into the experiment (Fig. 2B and 2C).

**Figure 2.**
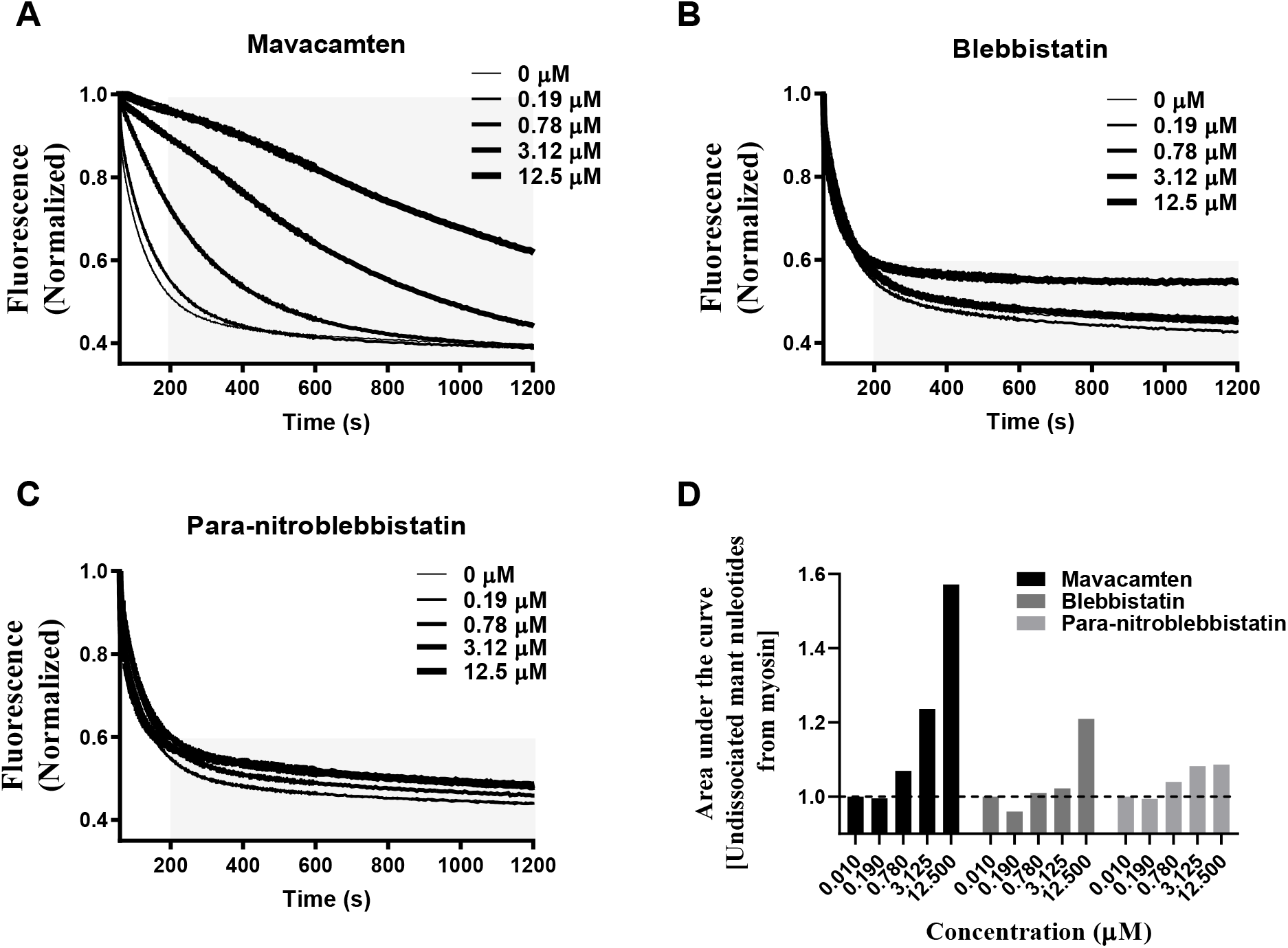
Mavacamten, but not Blebbistatin or Para-nitroblebbistatin, Slows the Release of Mant-Nucleotides from Myosin, in a Concentration-dependent Manner. Representative single turnover fluorescence decay profiles for (**A**) Mavacamten (**B**) Blebbistatin and (**C**) Para-nitroblebbistatin. (**D**) The area under the curve, which qualitatively describes the undissociated mant-nucleotides from myosin in a single turnover, is plotted for the traces shown in panels A-C. Only mavacamten shows a concentration-dependent increase in the undissociated mant nucleotide, while this is not the case with either blebbistatin or para-nitroblebbistatin below 12.5μM, respectively. The grey shaded region in panel A-C highlights the differences in fluorescence decay profiles from 200 sec onwards for various treatments.

The dose-dependent profiles of Aslow, as derived from a bi-exponential fit to these kinetic traces (see Methods and as described in (39)), which represent the % myosin population in the SRX state, are shown in Fig. 3A. The relative SRX population increased gradually with increasing mavacamten concentration, reaching 100% at 3.25 μM, suggesting that mavacamten shifted the myosin DRX-SRX equilibrium more towards the SRX state, consistent with what has been observed earlier (23, 25, 26). The concentration of mavacamten required to attain a half-maximal increase of myosin population in the SRX state is 1.21±0.35 μM, close to the IC_50_ of ATPase data measured in various systems (Table 1). In contrast, neither blebbistatin nor para-nitroblebbistatin showed any effect on Aslow up to a concentration of 12.5μM, indicating that neither of these two molecules affects the normal myosin DRX-SRX equilibrium. Qualitatively similar results were also obtained using skinned bovine cardiac myofibrils (Fig. 3C), verifying that the functional outcomes of these small-molecules are not different between a reconstituted thick filament system and a more intact myofibrillar system. Additionally, in both BcSTF (Fig. 3B) and BcMF (Fig. 3D), the ATP turnover rate of myosin in the DRX state (*k*_fast_) decreased in a concentration-dependent manner only for mavacamten but not for blebbistatin or para-nitroblebbistatin. This reinforces the notion that only mavacamten sequesters more myosin from the fast DRX state (with a rate of ~0.02 s^-1^) to the usual slow SRX state (with a rate of ~0.002 s^-1^) with increasing concentration. The concentration of mavacamten required to attain a half-maximal decrease (IC_50_) in *k*_fast_ is 0.43±0.03 μM, which is in good agreement with the IC_50_ values of basal, actin-activated, and myofibrillar ATPase activity (Table 1).

**Figure 3.**
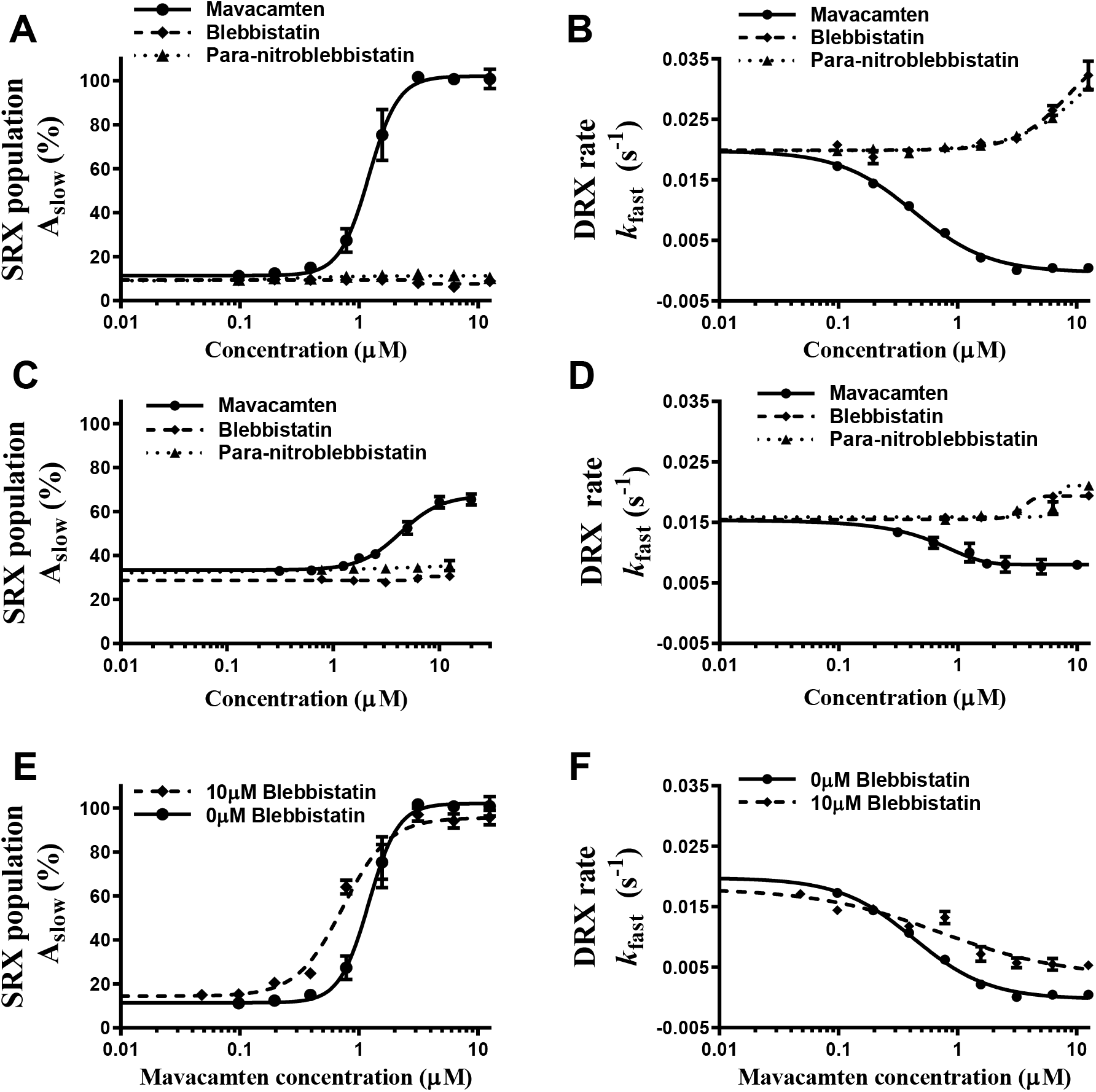
Mavacamten, but not Blebbistatin or Para-nitroblebbistatin, Populates Myosin in the SRX State. (**A**and **B**) Show comparison of the responses of mavacamten, blebbistatin, and para-nitroblebbistatin in BcSTF for SRX population (Aslow) and DRX rate (*k*_fast_), respectively, while (**C**and **D**) show respective comparisons in BcMF. (**E**and **F**) Compare the responses of mavacamten in BcSTF in the presence and absence of 10 μM blebbistatin for Aslow and *k*_fast_, respectively. BcSTF refers to bovine cardiac synthetic thick filaments, and BcMF refers to bovine cardiac myofibrils. Circles, diamonds, and triangles correspond to mavacamten, blebbistatin, and para-nitroblebbistatin, respectively. In either panel, the data were presented on the absolute scale. The half-maximal change (EC50/IC_50_) of mavacamten estimated from these curves are shown in Table 1. Data are expressed as mean±SEM (n≥8 from at least two experiments).

In another experiment, to determine whether the ability of mavacamten to increase the myosin SRX state is affected by the presence of blebbistatin, we performed single turnover experiments by pre-incubating BcSTF with a near-saturating concentration (10 μM) of blebbistatin and then titrating it with increasing concentrations of mavacamten. In the presence of 10 μM blebbistatin, the concentrations of mavacamten required to attain a half-maximal increase in myosin SRX state and half-maximal decrease in myosin DRX rate were 0.63±0.06 and 0.64±0.12 μM, respectively, which compare very well with those measured in the absence of blebbistatin (1.21±0.35 and 0.43±0.03 μM, respectively). Separately, we also confirmed that the binding of blebbistatin is not compromised in the presence of mavacamten (Fig. S1), suggesting that these two molecules do not compete for the same site, and the presence of blebbistatin does not affect the ability of mavacamten to populate more myosins in the SRX state.

### Mavacamten Stabilizes a More Compact Structure of Myosin Than Does Blebbistatin

To evaluate the effect of mavacamten and para-nitroblebbistatin on myosin structure, we used Small-Angle X-ray Scattering (SAXS) (Fig. S2) combined with Multi-Angle Light Scattering (MALS) and Dynamic Light Scattering (DLS) with purified human β-cardiac 25-hep heavy meromyosin (HMM) subfragment of myosin. The 25-hep HMM domain of the myosin used in this study consists of the two S1 domains and 25 heptads (25 repeated patterns of seven amino acids) of the S2 domain. Molecular weights estimated by MALS ensured the mono-dispersity of the sample. They were estimated to be 365(± 5%) kDa, 363(± 4%) kDa, 366(± 5%) kDa for DMSO-treated, para-nitroblebbistatin-treated, and mavacamten-treated 25-hep HMM, respectively (Table 1), which compares well to the theoretical molecular weight of ~380 kDa. These values were reproducible in 3 different repetitions within the experimental error and varied between 360 and 377 kDa. The hydrodynamic radius estimated by DLS for the 25-hep HMM was ~84 Å, with no substantial changes observed with either mavacamten or para-nitroblebbistatin (Table 1), suggesting that the apparent size adopted by the solvated, tumbling HMM does not change with either treatment. These studies were done with either 20 μM mavacamten or 50 μM para-nitroblebbistatin to ensure that we study their structural properties at complete biochemical inhibition levels (Fig. 1).

For isotropic solution scattering, scattering data reduction refers to the process of converting counts on the SAXS detector to the one-dimensional scattered intensity profile arising from the sample, with associated errors, as I(q) vs. q (where 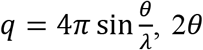 is the scattering angle and λ is the wavelength of the radiation). Guinier analysis (ln I(q) vs. q^2^) of the I(q) vs. q curves showed a significantly reduced radius of gyration (R_g_) for the mavacamten-treated HMM compared to para-nitroblebbistatin-treated and untreated 25-hep HMM (Fig. S3 and Table 1). The R_g_ values for the untreated, para-nitroblebbistatin-treated, and mavacamten-treated myosins were 119±1.6 Å, 101±1.3 Å, and 97±1.6 Å, respectively. The dimensionless Kratky plot ((qR_g_)^2^I(q)/I(0)versus qR_g_) is commonly used for qualitative assessment of flexibility and compactness of macromolecules. The relative downward shift of the Kratky plot for para-nitroblebbistatin-treated and mavacamten-treated HMM compared to that for DMSO-treated HMM indicated that the HMM molecule becomes more compact in the presence of either para-nitroblebbistatin or mavacamten, as compared to the untreated HMM (Fig. 4A).

**Figure 4.**
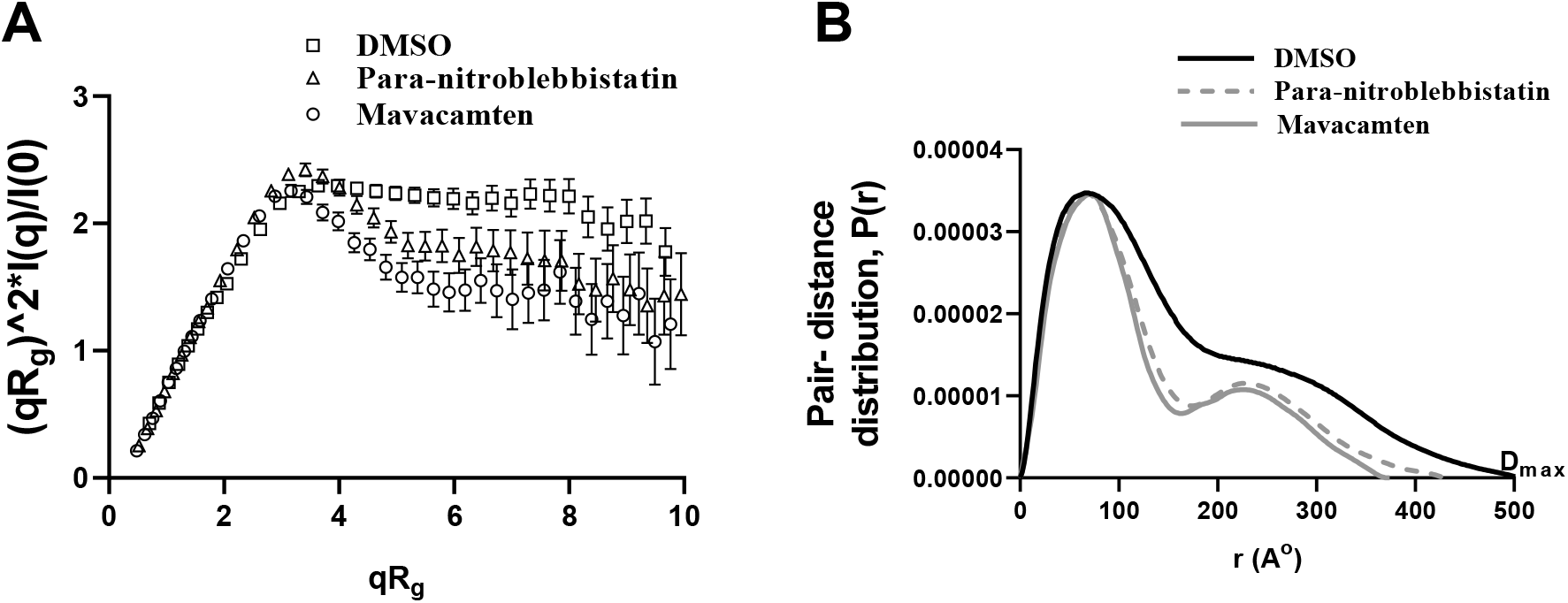
Mavacamten Stabilizes a More Compact Structure of Myosin as Compared to Blebbistatin. (**A**) Comparison of dimensionless Kratky plots from 25-hep HMM treated with DMSO, para-nitroblebbistatin, and mavacamten. Kratky plots are commonly used for qualitative assessment of flexibility and compactness of biological molecules and are independent of the molecular size. The downward shift of the Kratky plot indicates that 25-hep HMM becomes less flexible, probably more compacted, in the presence of either mavacamten or para-nitroblebbistatin as compared to control (DMSO). (**B**) Comparison of the pair-distance distribution function, P(*r*), of 25-hep HMM treated with DMSO, para-nitroblebbistatin, and mavacamten. P(*r*) functions of 25-hep HMM indicates an elongated structure with two globular domains consistent with the classic HMM structure. HMM treated with mavacamten and para-nitroblebbistatin elicited a decrease in the maximum dimension (D_max_) by different amounts compared to the untreated (DMSO) control. D_max_ is the maximum dimension of the molecule. At least 3 separate experiments were performed on different batches of samples for each treatment to confirm the reproducibility of the trend. Presented here are single representatives, which also produced the best fits during rigid body modeling shown in Fig. 5.

The pair-distance distribution function, P(*r*), which represents the frequency histogram of all the interatomic vector lengths within the sample particle, can provide structural information concerning the underlying distribution of electron density in the molecules. The P(*r*) of para-nitroblebbistatin-treated and mavacamten-treated 25-hep HMM showed similar features as the DMSO-treated HMM control (Fig. 4B). However, the maximum dimension (D_max_) of 25-hep HMM was reduced to a greater extent in mavacamten-treated samples than in para-nitroblebbistatin-treated samples. The D_max_ values for the untreated, para-nitroblebbistatin-treated, and mavacamten-treated myosins were ~ 505 Å, 430 Å, 380 Å, respectively (Table 1). Similarly, R_g_ values from the P(*r*) analysis of untreated, para-nitroblebbistatin-treated, and mavacamten-treated 25-hep HMM were 135±1.3 Å, 114±1.6 Å, and 106±1.4 Å, respectively (Table 1). These results collectively demonstrate that mavacamten induces greater compaction of myosin than does para-nitroblebbistatin, suggesting that these two molecules affect the structure of myosin differently.

### Mavacamten Populates a More Compact Closed State of Myosin than Para-nitroblebbistatin

To reconcile the SAXS data with known structural conformations, we attempted to fit the two starting human β-cardiac myosin homology models representing the hypothetical *open* and *closed* states as described in Nag et al. (9) (MS03, open-source material downloaded from http://spudlab.stanford.edu/homology-models) against SAXS data obtained from DMSO, para-nitroblebbistatin, and mavacamten-treated HMM using CRYSOL (40). *Open* and *closed* head conformations were modified by extending the tail and adding Green Fluorescent Protein (GFP, PDB-ID 4KW9) to the tail to match the 25-hep HMM construct used for the SAXS studies (Fig. 5A and B). SAXS curves representing untreated, para-nitroblebbistatin- and mavacamten-treated HMM fitted relatively well with the closed head conformation, when compared to the open head conformation (indicated by the χ^2^ values of 3.34, 3.17, and 2.61 respectively, as opposed to 20.56, 22.39 and 20.29; Fig 5A and B). Previous EM studies (23) and the failure to fit the *open* configuration suggest that DMSO-treated HMM, in solution, may be represented by either multiple conformations or a single alternate conformation. While the HMM treated with para-nitroblebbistatin or mavacamten seemed to have an obvious preference for the *closed* head conformation, the differences in R_g_ and D_max_ values from the Guinier and pair distance distribution analyses strongly indicated conformational differences, which suggested the need to optimize further the structural models resulting from small molecule treatment of HMM.

**Figure 5.**
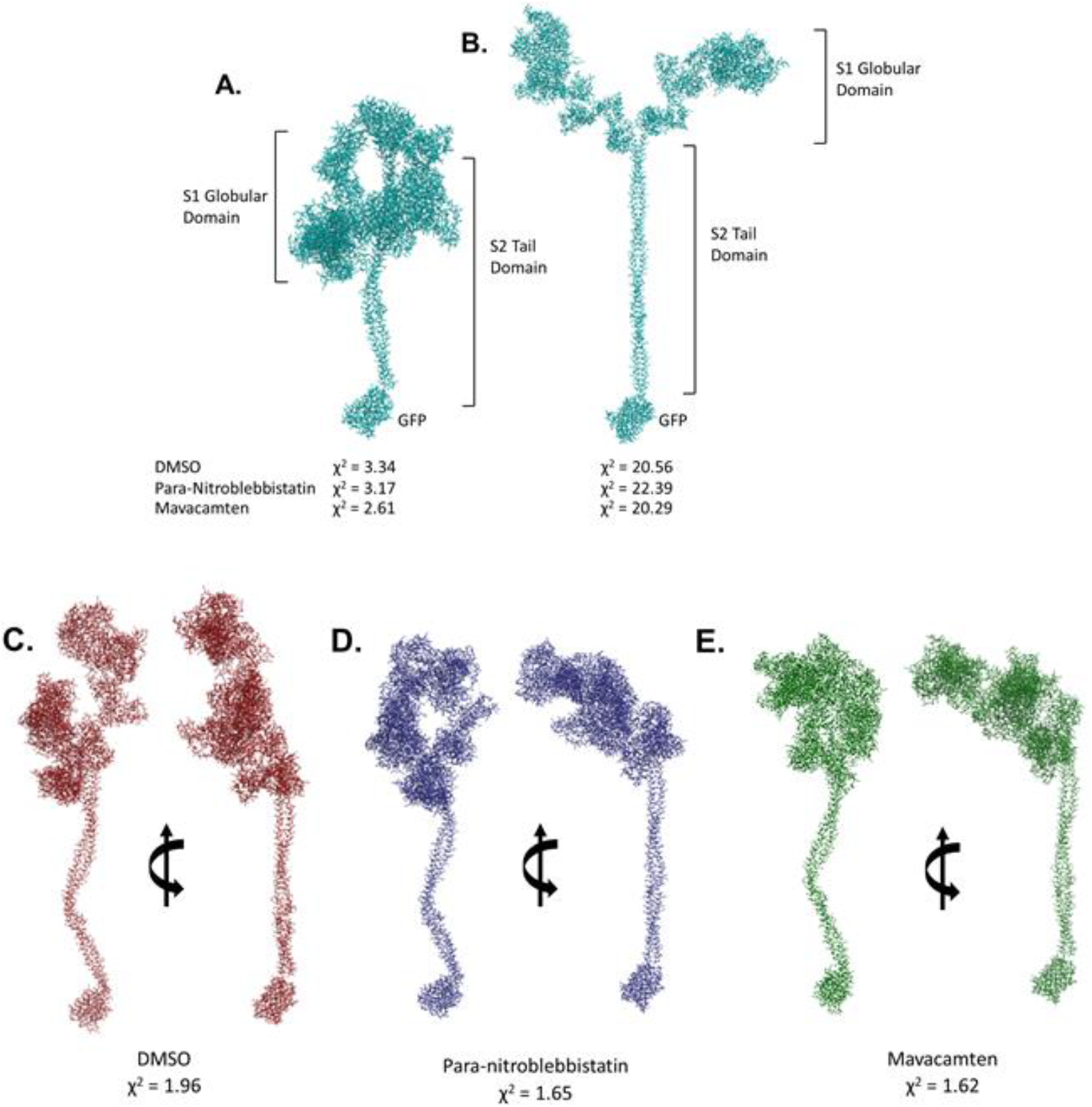
Rigid-body Modelling Using the SAXS data Suggests Conformational Heterogeneity and a Higher Propensity for Conformations Closer To the Closed Head Configuration in Mavacamten-treated HMM. Closed head **(A)** open head (**B**) conformation with the 25-hep tail and the GFP used for fitting against SAXS data obtained on untreated (DMSO), para-nitroblebbistatin, and mavacamten treated HMM using CRYSOL (40). χ^2^ values from CRYSOL indicated. Rigid body models obtained using SASREF (41) performed using open and closed head conformations, respectively, as starting models for HMM treated with (**C**) DMSO, (**D**) Para-nitroblebbistatin, and (**E**) Mavacamten. χ^2^ values from SASREF analysis are presented under the corresponding models.

We next used the rigid-body modeling software SASREF (41) to explore the possibility of a single conformation other than the hypothetical *open* and *closed* head starting models. We used SASREF to sample different relative orientations between the lever arm in the S1-subfragment and the proximal S2-subfragment domain in the hypothetical *closed* state because the initial CRYSOL fit was better for all three preparations compared to the *open* state. The S1-S2 disordered hinge is known to have flexibility in all myosins providing the S1 heads the needed rotational freedom to make productive interactions with actin (recently illustrated in the review (42)). Multiple SAXS experiments were performed on different preparations of HMM, and numerous iterations of SASREF were performed with the *closed* conformation as a starting model. DMSO-treated HMM converged to a more extended conformation than the theoretical *open* conformation and was consistent with the higher D_max_ and R_g_ estimates (Fig. 5C). Para-nitroblebbistatin-treated and mavacamten-treated HMM consistently reached a conformation best described as an intermediate between the *open* and *closed* states (Fig. 5D and E). While the P(r) analysis showed a difference in D_max_ between para-nitroblebbistatin-treated and mavacamten-treated HMM, SAXS-based rigid body modeling proved to be inadequate for distinguishing the specific structural features between the two. Based on these data, we posit that while para-nitroblebbistatin and mavacamten both induce conformations between the hypothetical *open* and *closed* states, the average myosin structure stabilized by mavacamten is more compact than by para-nitroblebbistatin. We also note that it is quite likely that the ensemble structure that myosin adapts in the presence of these agents may not be well-represented by any linear interpolation of the “canonical” structures. Notably, the *open* conformation is probably not a single configuration, but instead, myosin can undergo many structural reconfigurations as it advances through the chemomechanical cycle.

## DISCUSSION

The use of small-molecule agents that target cardiac myosin and inhibit its ATPase activity is emerging as an attractive therapeutic strategy to normalize excessive contractility in hypertrophic cardiomyopathy. In this regard, although several clinical and non-clinical small molecule inhibitors have been identified and characterized, the underlying mechanism of their inhibitory actions remains unclear. This study aimed at filling some of this gap by investigating the structure-function relationships of cardiac myosin using three small-molecule inhibitors—mavacamten, blebbistatin, and para-nitroblebbistatin. It has been known that mavacamten and blebbistatin are myosin inhibitors, although much of the data on the inhibitory action of blebbistatin comes from studies using skeletal muscle, smooth muscle, and non-muscle myosin systems (27–29). Here, we found that, in the presence of mavacamten, blebbistatin, and para-nitroblebbistatin, the inhibition of basal myosin ATPase activity was comparable to the inhibition observed in the cardiac actomyosin or myofibrillar system (Fig. 1 and Table 1), suggesting that all the three molecules directly acted upon myosin to inhibit its enzymatic activity and that this effect does not involve any other sarcomere proteins, including the regulatory components of the thin filaments. This leads to an immediate question “how do these small molecules inhibit cardiac myosin function?”

Analysis from single turnover experiments showed that mavacamten, but not blebbistatin or para-nitroblebbistatin, decreased the myosin population in the DRX state that can readily interact with actin by consequently increasing the population in the energy-sparing SRX state. This explains both the inhibition in ATPase activity (Table 1) and the observed reduction in cardiac contractility in pre-clinical and clinical models (21, 43–49). This idea is also qualitatively substantiated by the regression analysis, which demonstrates that the inhibition in either the basal or the actin-activated myosin ATPase activity by mavacamten linearly correlates to the increase in myosin SRX population (Fig. 6). In contrast, at inhibiting concentrations, neither blebbistatin nor para-nitroblebbistatin altered the myosin SRX state (Fig. 3 and 6). This finding agrees with another study using rabbit slow skeletal muscle fibers that expresses the same myosin isoform as cardiac muscle (37). Specifically, the authors showed that, at a saturating concentration of 40 μM, blebbistatin does not affect the early fluorescence decay phase in the single turnover experiments but considerably slows the rate of the late phase, which corresponds to the myosin SRX state. This observation could possibly be explained by blebbistatin stabilizing a small but significant fraction of SRX heads into a third ‘ultra-relaxed’ state in which myosins have greater than 10-fold lower ATP turnover rate than those in the usual SRX state, thereby practically manifesting these myosins to have negligible ATPase function (Fig. 1). Our data at low concentrations (Fig. 2) and a high blebbistatin concentration of 50 μM (Fig. S4) (close to 40 μM used in the previous study) supports such a possibility. Mavacamten and blebbistatin populating different relaxed states is further supported by our observation that mavacamten can populate myosin heads into the normal SRX state with similar effectiveness even in the presence of blebbistatin bound to myosin, suggesting that blebbistatin binding to myosin does not alter the usual DRX-SRX equilibrium of myosin. Altogether, these observations suggest the possibility that unlike mavacamten, binding of blebbistatin, near the nucleotide-binding site of myosin (50), but far from the myosin lever arm, renders myosin to unable to prime the lever arm more towards the pre-stroke direction, which is proposed to be a pre-requirement to populate the normal SRX state (14, 18).

**Figure 6.**
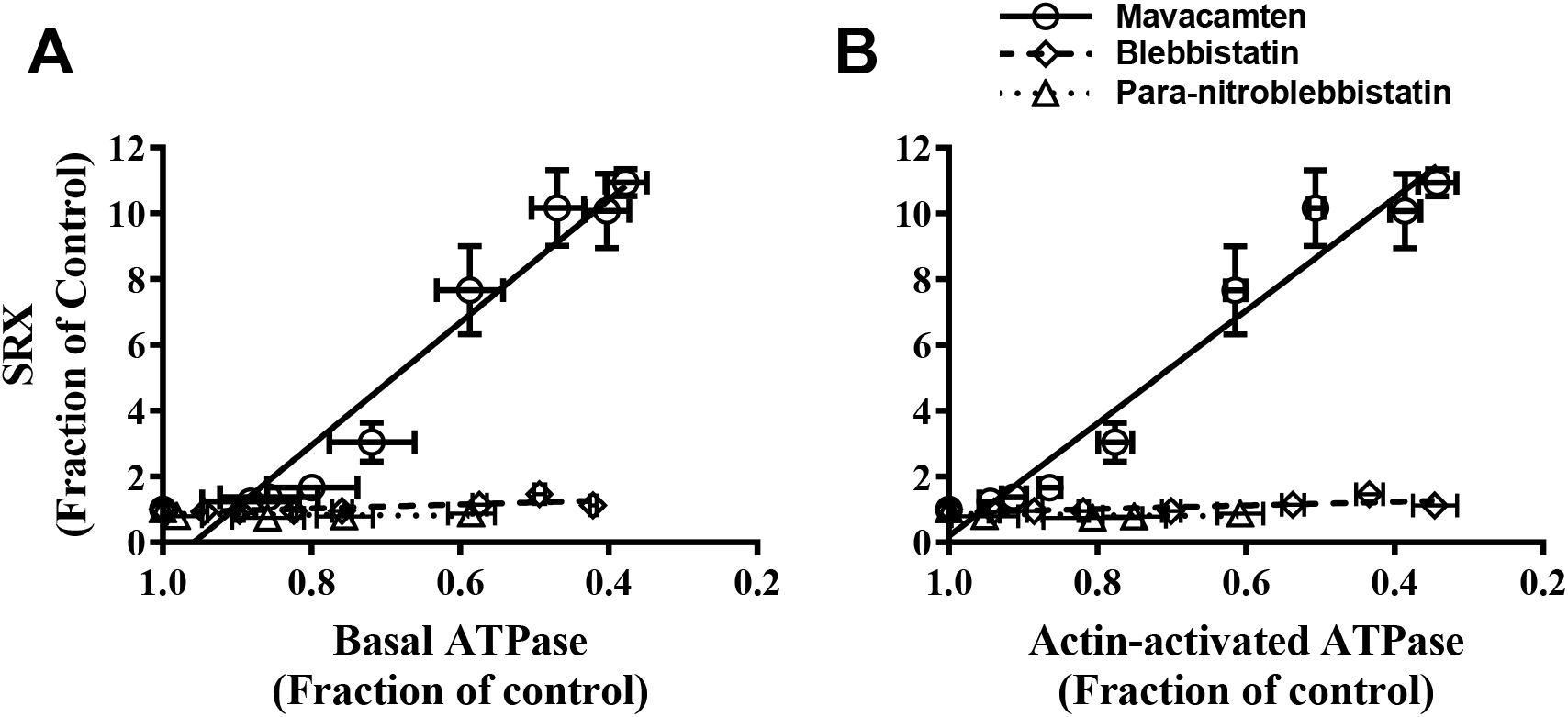
Inhibition of Myosin ATPase Caused by Mavacamten, but not Blebbistatin or Para-Nitroblebbistatin, Linearly Correlates with Increase in Myosin SRX Population. Correlation plot between myosin SRX population and (**A**) basal myosin ATPase activity and (**B**) actin-activated ATPase activity for mavacamten (circles), blebbistatin (diamonds), and para-nitroblebbistatin (triangles). Regression analysis showed a strong linear correlation between myosin population in SRX state and ATPase activity for mavacamten (R^2^≥0.94) but not for blebbistatin or para-nitroblebbistatin. All data were normalized by the DMSO-treated sample data and presented as mean±SEM (*n*≥8 for each condition).

Additional evidence that para-nitroblebbistatin structurally affects myosin differently from mavacamten comes from structural parameters derived using SAXS/MALS/DLS measurements. For example, while the Kratky plots suggested that both mavacamten and para-nitroblebbistatin qualitatively decrease the flexibility and increase the compactness of the 25-hep HMM, parameters derived from pair-distance distribution function (P(*r*)) quantified the differences among treatments. In the P(*r*) analysis, the DMSO-treated HMM showed a bell-shaped curve with a peak at low *r* values (0-170Å), a shoulder at intermediate *r* values (170-340Å), and an extended tail at high *r* values (>340Å), indicating an extended structure with D_max_ of ~ 505 Å (Fig. 4B). Intuitively, this kind of P(*r*) functions for proteins represent a long-tailed structure with globular domains, consistent with the classic HMM structure in which the S2-subfragment of two myosin molecules form a coiled-coil rod on which the flexible globular S1 heads rests. The bell-shaped curve in P(*r*) at low *r* values presumably comes from the rigid, well-defined intra-head vector length, while the broad shoulder in P(*r*) for intermediate *r* values comes mostly from the inter-head vector lengths indicating a flexible inter-head spacing. Interestingly, compared to untreated (DMSO) and para-nitroblebbistatin-treated samples, the shoulder region of P(*r*) for mavacamten-treated HMM elicited a more prominent peak with higher probability, indicating a qualitatively well-defined, less scattered inter-head spacing. Significantly lower values for physical parameters such as R_g_ and D_max_ in mavacamten-treated HMM compared to both para-nitroblebbistatin-treated HMM and the DMSO-treated control (Table 1) strongly suggest that mavacamten induces a much more compact and rigid form of myosin than para-nitroblebbistatin, supporting the idea that mavacamten structurally populates myosin in a different *closed* state when compared to para-nitroblebbistatin. To strengthen our assertion, rigid body modeling using SASREF (41) converged to a myosin conformation that was more extended than the theoretical open conformation (Fig. 5C), which when paired with a broader shoulder region in the P(*r*) distribution (Fig. 4) and higher ensemble averages of R_g_ and D_max_ (Table 1) suggest that there is a preferential shift in the distribution of myosin more towards the *open* state in the DMSO-treated HMM. However, in the HMM samples treated with either mavacamten or para-nitroblebbistatin, the SASREF modeling converged to a single homogeneous conformation that may be best described as a structural intermediate between the *closed* and *open* states (Fig. 5D–E) but proved to be inadequate to distinguish the structural features between treatments. However, these modeling-based solutions when combined with other structural parameters such as a narrow shoulder region in the P(*r*) distribution indicating less inter-head flexibility, paired with lower R_g_ and D_max_, indicated that the degree of compactness in myosin is greater in the presence of mavacamten than with para-nitroblebbistatin and that mavacamten shifted the myosin distribution more towards a compact *closed* state.

It is worth noting that, in all HMM samples, including the control (DMSO), neither a classic *closed* myosin conformation mimicking the actual IHM state (i.e., two heads in the dimeric myosin folding back and interacting with the S2-subfragment tail) nor a genuine *open* conformation best described the model predicted by the SASREF modeling. While these findings differ from those of the previous studies, which show that both mavacamten and blebbistatin can promote myosin into a fully compact folded-state (23, 33), they are consistent with other findings from earlier reports. For example, findings from an X-ray diffraction study show that mavacamten treatment increases myosin density along thick filaments in cardiac muscle fibers, reflecting a compact arrangement (23). Likewise, electron microscopy and structural studies using fluorescent polarization show that blebbistatin induces compactness by aligning myosin heads closer and parallel to the thick filament backbone (36) and improves the helical ordering of myosin along thick filaments in skeletal systems (31, 32). The discrepancy between findings from this study and previous studies may be likely attributed to the differences in the techniques used (e.g., solution-based SAXS structure vs. static electron microscopy images). Also, hydrolysis of ATP to different nucleotide states in solution during the chemomechanical cycle can render myosin into different conformations, and the modeling of SAXS data presents us with an ensemble average of these structures. Also, mavacamten and blebbistatin may not bind to myosin equally in all these nucleotide states, thereby giving different weights to different structures in the ensemble averaging.

Collectively, these studies lead us to posit that blebbistatin (or para-nitroblebbistatin) is a “poison” that renders myosin in a structural intermediate between *closed* and *open* states that prevents myosin from undergoing a productive ATPase cycle by probably populating an ‘ultra-relaxed’ state, which dramatically inhibits nucleotide release. This mechanism needs to be fully uncovered using better approaches in future investigations. In contrast, mavacamten shifts the *open*-*closed* equilibrium more towards the *closed* state and inhibits myosin ATPase function by increasing the energy-sparing myosin SRX population by most likely promoting myosin into more compact conformational states in which the lever arm may be primed towards the pre-stroke direction(14, 23). Inferences drawn from our study support the idea that, in addition to the classic IHM state, different interventions may structurally stabilize myosin in many other *closed* states, and not all such *closed* states may necessarily correlate to the functional SRX state. Our study provides novel insights into the mechanistic actions of small molecules by connecting structural properties to biological functions, which is fundamental to designing better molecules for improved therapies.

## MATERIALS AND METHODS

All materials and methods are briefly described here, but more details can be found in the Supporting Information (SI).

### Preparation of Protein Reagents

Bovine cardiac actin, bovine cardiac full-length myosin, and bovine cardiac myofibrils were prepared following the methods described previously (51–53). Human β-cardiac 25-hep heavy meromyosin (HMM) was purified using methods described elsewhere (54). The human β-cardiac 25-hep HMM cDNA consists of a truncated version of MYH7 (residues 1-855), corresponding to S1-subfragment and the first 25 heptad repeats (175 amino acids) of S2-subfragment, followed by a GCN4 leucine zipper to ensure dimerization. This is further linked to a flexible GSG (Gly-Ser-Gly) linker, then a GFP moiety followed by another GSG linker, and finally ending with an 8-residue (RGSIDTWV) PDZ binding peptide.

### Reconstitution of Myosin Thick Filaments

Starting from a buffer of high ionic strength (300 mM KCl or higher) in which full-length myosin remains completely soluble, the buffer’s ionic strength was reduced to 30 mM to allow spontaneous self-assembly of bi-polar thick filaments as described previously (55, 56). In this study, bovine β-cardiac full-length myosin has been used to form such reconstituted myosin filaments and are referred to as bovine cardiac synthetic thick filaments (BcSTF).

### Steady-state ATPase Measurements

Measurements of basal (pCa 10), actin-activated, and myofibrillar (pCa 6) myosin ATPase activity were performed at 23°C on a plate-based reader (SpectraMax 96-well) using an enzymatically coupled assay described elsewhere (21, 38). The buffer conditions used were 12 mM Pipes (pH 6.8), 2 mM MgCl2, 10 mM KCl, and 1 mM DTT. All basal and actin-activated ATPase experiments were done with myosin in the form of STFs. A final myosin concentration of 1 μM was attained in all cases. The concentration of actin in the actin-activated myosin ATPase experiments was 14 μM. When using myofibrils, approximately 40% of the total weight was assumed to be from myosin, and accordingly, the myofibril amount was loaded to attain a final myosin concentration of 1 μM.

### Preparation of Myosin-based Small Molecule Inhibitors

Three different small molecule myosin inhibitors, blebbistatin, para-nitroblebbistatin, and mavacamten, were used in this study. Mavacamten was synthesized by MyoKardia, Inc., whereas blebbistatin and para-nitroblebbistatin were purchased from external sources (blebbistatin from Fisher Scientific, IL, and para-nitroblebbistatin from Cayman Chemical Company, MI). Concentrated stocks (20 mM) of each molecule were first prepared using dimethyl sulfoxide (DMSO). These are used to achieve a final concentration ranging from 0 to 50 μM in the experimental buffer samples. The final concentration of DMSO was 2% in all cases. Structural studies involving Small-angle X-ray scattering (SAXS), Multi-Angle Light Scattering (MALS), and Dynamic Light Scattering (DLS) measurements were carried out using a saturating concentration of each compound (20 μM for mavacamten and 50 μM for para-nitroblebbistatin).

### Single nucleotide Turnover Experiments

Single ATP turnover kinetic experiments using a fluorescent 2′/3′-O-(N-Methylanthraniloyl) (mant)-ATP were conducted in a 96-well plate fluorescence plate reader, as described in detail in previous work (26). Briefly, in the first step, 50 μL of the experimental buffer containing 3.2 μM mant-ATP is combined with 100 μL of the same buffer containing 0.8 μM myosin in a UV plate. The reaction was aged for 60 s to allow binding and hydrolysis of mant-ATP to inorganic phosphate (Pi) and mant-ADP. In the second step, mant-nucleotides were chased with non-fluorescent ATP by adding 50 μL of 16 mM non-fluorescent ATP to the above mixture, and the resulting fluorescence decay due to mant-nucleotide dissociation from myosin was monitored over time. The experimental buffer contained the following: 20 mM Tris-HCl (pH 7.4), 30 mM KCl, 1 mM EGTA, 3 mM MgCl2, and 1 mM DTT. The concentrations of myosin, mant-ATP, and non-fluorescent ATP attained in the final mixture were 0.4 μM, 0.8 μM, and 4 mM, respectively. The excitation wavelength for mant-nucleotides was 385 nm. The emission was monitored using a long-pass filter with a cutoff wavelength of 465 nm. All experiments were performed at 25°C.

As described in other studies (23, 25, 39), the fluorescence decay profile during the chase phase characteristically depicted two phases, a fast phase followed by a slow phase. Therefore, a bi-exponential function was fitted to each trace to estimate four parameters corresponding to fast and slow phases — Afast, *k*_fast_, Aslow, and *k*slow — where A represents the % amplitude, and *k* represents the observed ATP turnover rate of each phase. In this study, the fast and slow phases correspond to myosin activity in the DRX and SRX states, respectively.

### SEC-MALS-DLS-SAXS Studies

Small-angle X-ray scattering (SAXS) was performed with in-line size-exclusion chromatography (SEC), multi-angle light scattering (MALS), and dynamic light scattering (DLS) at the BioCAT beamline 18ID (Advanced Photon Source, Argonne National Laboratory, Lemont IL). The 25-hep HMM was loaded on a superose-6 increase 10/300 column (GE Lifesciences), which was run at 0.7 ml/min on an Agilent Infinity II HPLC system. The column eluant passed through a UV detector, MALS and DLS detectors (Wyatt DAWN HELEOS II), and a differential refractive index detector (Wyatt Optilab t-rEX) before reaching a co-flow SAXS cell. Samples and experimental buffers were treated with DMSO, 20 μM mavacamten, or 50 μM para-nitroblebbistatin. Because these experiments were run for a longer period (~2 hours), we avoided using blebbistatin in these experiments due to its unstable nature in the aqueous medium (57). Instead we used the non-cytotoxic and photostable para-nitroblebbistatin derivative (58). All such experiments were done at 25°C, and the buffer used for these experiments contained 20 mM Tris-HCl (pH 7.4), 30 mM KCl, 1 mM EGTA, 3 mM MgCl2, and 1 mM DTT.

Scattering intensity was recorded using a Pilatus3 1M detector, which was placed ~3.5m from the sample giving access to a q range of ~0.004Å^-1^ to 0.4 Å^-1^. 0.5-second exposures were acquired every second during elution, and data were reduced using BioXTAS RAW 1.6.0 (59). Buffer blanks were created by averaging regions flanking the elution peak and subtracted from exposures selected from the elution peak to create the I(q) vs. q curves used for all subsequent analyses, which included Guinier approximation (radius of gyration, R_g_ and zero-angle scattering, I0), pair-distance distribution function (R_g_ and the maximum dimension – D_max_) and calculation of structural models. ASTRA (Wyatt Inc.) was used to calculate the molecular weight and hydrodynamic radii based on the MALS and DLS data, respectively. Four repeat experiments were performed on four different preparations of HMM to confirm the trends in the calculated parameters, and the results presented here are from the experiments with the best signal to noise.

### Data Analysis

For biochemical studies, each assay was repeated at least twice with a minimum of two replicates per experiment. Sample number *n* refers to the total number of measurements made (number of experiments*number of replicates), and these data were presented as mean±SEM. The dose-response curve in each assay was used to estimate the concentration (in μM) required to attain half-maximal decrease (IC_50_) or half-maximal increase (EC50) in a given parameter for each small molecule treatment. Linear regression analysis was performed between data from various experiments to determine the strength of such correlations. For SAXS studies, experiments were repeated at least three times for each treatment (DMSO, para-nitroblebbistatin, mavacamten), and representative data (lowest chi-squared values for fits) are presented hereafter confirming the reproducibility of the fundamental trend.

## DATA AVAILABILITY

All data presented in this manuscript is either included in the main article or the Supporting Information.

## AUTHOR CONTRIBUTIONS

SKG and SN conceptualized the study design and formulated the hypotheses. SKG designed, collected, analyzed, and interpreted all biochemical data. WM and SC designed, collected, analyzed, and interpreted all SAXS data. ACC and SL designed and constructed the recombinant 25-hep human HMM construct. ACC purified most of the 25-hep human HMM protein used for the SAXS studies. NS contributed to the initial preparations of the human 25-hep HMM and the biochemical ATPase data. SKG, SN, WM, SC and TCI wrote the manuscript and finalized the final contents. SN supervised the entire study.

## ACKNOWLEDGMENTS

The authors would like to thank the protein production team and the medicinal chemistry team of MyoKardia Inc. for the production of biological reagents and mavacamten. The authors thank Dr. Leslie Leinwand at the University of Colorado, Boulder, for providing supervision to produce cell pellets and purifying the human β-cardiac 25-hep HMM. The authors also thank Dr. Roger Cooke, Professor Emeritus at UCSF; Dr. Robert McDowell, Chief Scientific Officer of MyoKardia Inc.; and Robert Anderson, Biology leader at MyoKardia Inc., for providing critical inputs to the manuscript. This research used resources of the Advanced Photon Source, a U.S. Department of Energy (DOE) Office of Science User Facility operated for the DOE Office of Science by Argonne National Laboratory under Contract No. DE-AC02-06CH11357. This project was supported by grant P41 GM103622 from the National Institute of General Medical Sciences of the National Institutes of Health. The content is solely the responsibility of the authors and does not necessarily reflect the official views of the National Institute of General Medical Sciences or the National Institutes of Health.

