## Supplemental information for "TWO CLASSES OF MYOSIN INHIBITORS, BLEBBISTATIN AND MAVACAMTEN, STABILIZE β-CARDIAC MYOSIN IN DIFFERENT STRUCTURAL AND FUNCTIONAL STATES"


**Running title**: Differential stabilization of cardiac myosin by mavacamten and blebbistatin.


**Keywords:** Super-Relaxed State (SRX), Interacting Heads Motif (IHM), Small Angle-X-ray Scattering (SAXS), mavacamten, blebbistatin, human β-cardiac myosin, Synthetic thick filaments (STF)

### Human β–Cardiac HMM Preparations

The human β-cardiac 25-hep HMM construct was produced using a modified AdEasy™ Vector System (Qbiogene, Inc). The human β-cardiac 25-hep HMM cDNA consists of a truncated version of MYH7 (residues 1-855), corresponding to S1, and the first 25 heptad repeats (175 amino acids) of the myosin S2 -subfragment, followed by a GCN4 leucine zipper to ensure dimerization. This is further linked to a flexible GSG (Gly-Ser-Gly) linker, then a GFP moiety followed by another GSG linker, and finally ending with an 8-residue (RGSIDTWV) PDZ binding peptide. The recombinant 25-hep HMM construct was co-expressed with a FLAG-tagged human ventricular essential light chain (ELC) construct in C2C12 mouse myoblast cells using adenoviral vectors. Complete cloning, expression, and purification methodology are described in detail elsewhere (1, 2). Briefly, cDNA for the human cardiac myosin heavy chain (MYH7) and human ventricular ELC (MYL3) were purchased from Open Biosystems (Thermo Scientific, Waltham, MA). The human ventricular ELC with an N-terminal FLAG tag (DYKDDDDK) and TEV protease site was co-expressed with the heavy chain using an adenoviral vector/mouse myoblast C2C12 system (purchased from ATCC), as described previously (1). Purified fractions of the protein were either dialyzed or buffer-exchanged using Amicon centrifugal filter units (Millipore, Burlington, MA) in binding buffer containing 20 mM Tris-HCl (pH 7.4), 30 mM KCl, 1 mM EGTA, 3 mM $MgCl_2$, and 1 mM DTT and 5% sucrose at pH 7.5. Before any experiment, the myosin constructs were sedimented by centrifugation at 350,000 × g for 15 minutes to remove any aggregated protein. Protein concentrations were quantified by measuring the GFP moiety's absorbance at 488 nm on a Nanodrop spectrophotometer (Thermo Scientific, Waltham, MA).

### Blebbistatin Binding to Myosin is not Affected by the Presence of Mavacamten

It is previously known that blebbistatin has fluorescence properties and that its fluorescence increases upon binding to myosin (3–5). We utilized this property of blebbistatin to study whether its binding to myosin is affected by the presence of mavacamten. In this experiment, we pre-incubated bovine cardiac synthetic thick filaments (BcSTF) with either DMSO or 10µM mavacamten and exposed them to progressively increasing concentrations of blebbistatin, and measured the steady-state fluorescence in these samples on a 96-well plate fluorescence plate reader. The excitation wavelength was set to 434 nm for these measurements, and the emission was monitored using a 550 nm cut-off filter. We also performed similar measurements without including myosin in all samples to determine the intrinsic fluorescence ascribed to the blebbistatin alone. We then deducted these fluorescence values from those measured in respective samples containing myosin to estimate the fluorescence increase ascribed to the blebbistatin binding to myosin. Comparison of such profiles illustrated in Fig. S1 demonstrates that the binding of blebbistatin to myosin is not affected by the presence of mavacamten but is indeed slightly enhanced at all concentrations of blebbistatin. The concentrations required for the half-maximal increase in the fluorescence of blebbistatin were 40.6±4.4 µM and 21.7±1.81 µM in the presence and absence of 10 µM mavacamten, respectively.

**SAXS/MALS-DLS Studies**

SAXS experiments were performed using the BioCAT beamline 18ID at the Advanced Photon Source, Argonne National Laboratory, Lemont, IL, with in-line size exclusion chromatography (SEC). The use of SEC ensured the separation of the sample from aggregates and other contaminants, thus ensuring optimal sample quality. All SAXS experiments were conducted using 25-hep human heavy meromyosin (HMM). In each experiment, 250 µL of HMM (3 mg/ml) treated with DMSO, 50 µM para-nitroblebbistatin or 20 µM mavacamten was loaded onto a Superose 6 Increase 10/300 GL column after 15 min centrifugation at 16000xg at 4°C. The matching elution buffer (20 mM Tris-HCl (pH 7.4), 30 mM KCl, 1 mM EGTA, 3 mM $MgCl_2$, and 1 mM DTT) was run at 0.6 ml/min by an Infinity II HPLC (Agilent Technologies, Santa Clara, CA). The sample to detector distance for the SAXS camera was ~3.5 m, which provided access to a q-range of ~0.004 to 0.4 $Å^{-1}$. Scattering intensity was recorded using a Pilatus 3 1M (Dectris) detector with a 0.5 s exposure for every second during elution. Before reaching the SAXS flow cell, the sample was sent to (in order) the Agilent Infinity II UV monitor MALS, DLS, and RI detectors (Wyatt Technologies, Santa Barabara, CA). MALS provided robust molecular weight estimates that corroborated the ones obtained by SAXS. Molecular weights and hydrodynamic radii were calculated from the MALS and DLS data using the ASTRA 7 software (Wyatt Technologies). More details on SAXS data collection and analysis are listed in Table S1.

The SAXS data were reduced using BioXTAS RAW 1.6.0 (6), and the summed scattering intensity was plotted against frame number to generate SAXS chromatographs or scattergrams (Figure S2; open symbols). Buffer blanks were created by averaging regions flanking the elution peak, and these are subtracted from exposures corresponding to the elution peak. The radius of gyration ($R_g$) value for each frame (Figure S2; solid symbols) was then calculated by BioXTAS RAW and plotted in the SAXS chromatograph. The frames corresponding to the protein peak were chosen and averaged by BioXTAS RAW automatically. One-dimensional SAXS curves were reduced from averaged frames by radial averaging for detailed analysis. The resulting I(q) vs. q curves were fit against hypothetical models which were generated by extending the 18-hep tail in pre-existing models (MS03, open-source material downloaded from http://spudlab.stanford.edu/homology-models) (2) to a 25-hep tail and also the addition of green fluorescent protein to the end of the 25-hep tail, all of which was accomplished using PYMOL (The Pymol Molecular Graphics System, Version 2.0 Schrodinger, LLC). Rigid body modeling was performed using SASREF (7), which uses simulated annealing to optimize the spatial arrangement of domains to fit the experimental data better. The model representing either the open or closed conformation was split into three separate domains, two head and neck regions and the tail with the GFP tag. Distance restraints between terminal residues of each domain were used to maintain sequence continuity and avoid steric clashes between the different domains. The models resulting from multiple runs of SASREF were chosen based on the best discrepancy value $\chi^{2,}$ which is defined by the equation –

$$\chi^2 = \frac{1}{N-1}\sum_j \left[\frac{I\exp(s_j) - cI(s_j)}{\sigma(s_j)}\right]^2$$

Where, $I_{exp}(s_j)$ is experimental scattering, $I(s_j)$ is the calculated scattering, N is the number of experimental points, c is a scaling factor, and $\sigma(s_j)$ is the experimental error at the momentum transfer $s_j$.

### SUPPLEMENTAL DATA TABLES

**Table S1.** SAXS Data Collection and Analysis Parameters

| | |
|---|---|
| a) SAXS data collection parameters | |
| *Instrument* | BioCAT facility at the Advanced Photon Source beamline 18ID with Pilatus3 1M (Dectris) detector |
| *Wavelength (Å)* | 1.033 |
| *Beam size* (μm$^2$) | 150 (h) x 80 (v) |
| Camera length (m) | 3.5 |
| q-measurement range (Å-1) | 0.0045-0.38 |
| Absolute scaling method | N/A |
| Basis for normalization to constant counts | To incident intensity, by ion chamber counter |
| Method for monitoring radiation damage | Automated frame-by-frame comparison of relevant regions |
| Exposure time, number of Exposures | 0.5 s exposure time with a 1 s total exposure period of entire SEC elution |
| Sample configuration | SEC-MALS-DLS-RI-SAXS. Size separation used a Superose 6 Increase 10/300 GL column and an Infinity II HPLC (Agilent Technologies). Flow was split between SAXS and the UV-MALS-DLS-RI instruments after the column. SAXS data was measured with sheath-flow cell (8) in a 1.5 mm ID 1.52 mm OD quartz capillary, effective path length 0.49mm. UV data was measured in the Agilent, and MALS-DLS-RI data by DAWN HELEOS-II (17 MALS + 1 DLS channels) and Optilab T-rEX (RI) instruments (Wyatt). |
| Sample temperature (ºC) | 22 |
| b) *Software for SAS data reduction, analysis, and interpretation* | |
| *SAXS data reduction* | Radial averaging, background subtraction, frame comparison, and frame averaging were made using BioXTAS RAW 1.6.0 **(6)**and ATSAS **(9)**. |
| *Basic analysis: Guinier, M.W., P(r)* | Guinier fit and molecular weights were calculated using BioXTAS RAW 1.6.0 **(6)**, P(r) function was estimated using GNOM **(9)**. |
| *MALS-DLS-RI analysis* | Astra 7.1.3 (Wyatt) |

**SUPPLEMENTAL FIGURES**

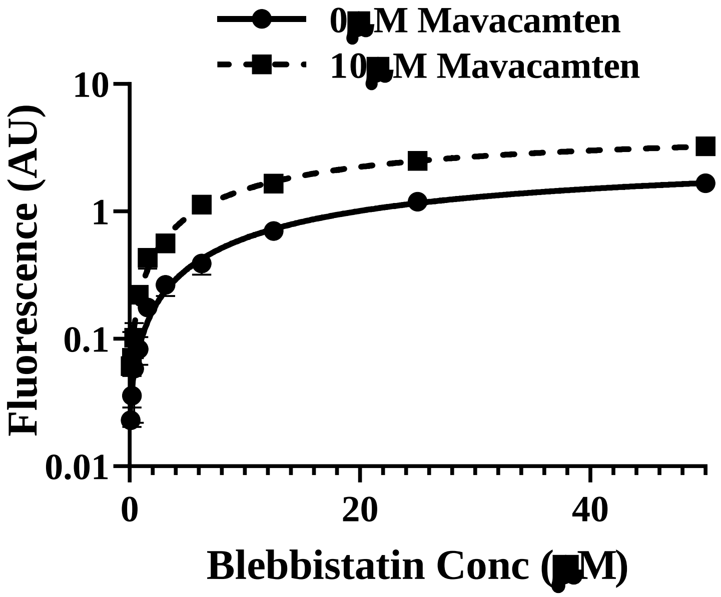


**Figure S1.** Fluorescence profiles of blebbistatin binding to BcSTF in the presence and absence of 10 µM mavacamten. The concentrations required for the half-maximal increase in the fluorescence of blebbistatin were 40.6±4.4 µM and 21.7±1.81 µM in the presence and absence of 10µM mavacamten, respectively.

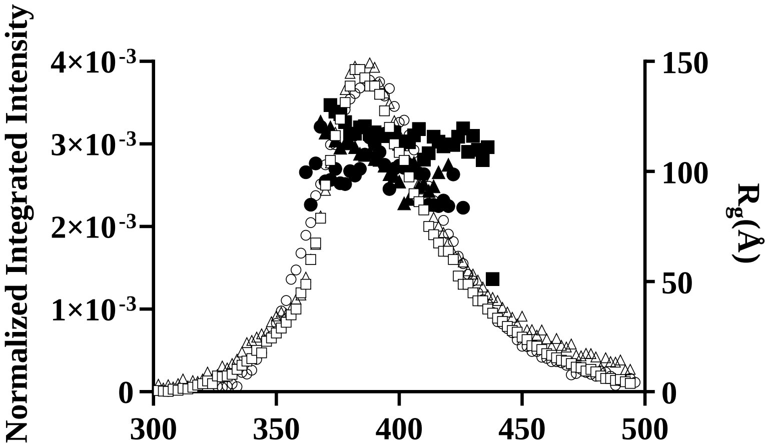

**Figure S2.** Representative SAXS scatterogram for human β-cardiac 25-hep HMM in the absence or presence of various small molecule inhibitors. Open symbols represent the normalized integrated intensity, while the solid symbols represent $R_g$ (DMSO: squares; para-nitroblebbistatin: triangles; mavacamten: circles).

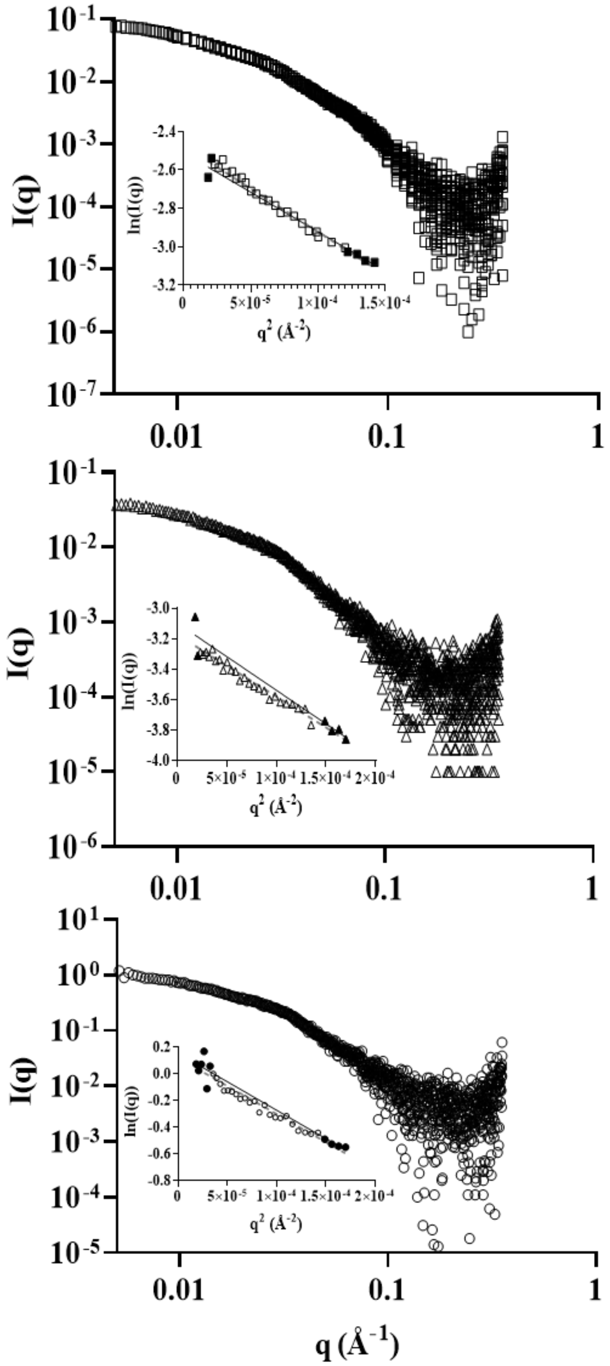


**Figure S3.** I(q) vs. q curves and Guinier plots (inset) for human β-cardiac 25-hep HMM in the presence and absence of small molecule inhibitors. Symbols correspond to the following: squares for DMSO; triangles for para-nitroblebbistatin; circles for mavacamten.

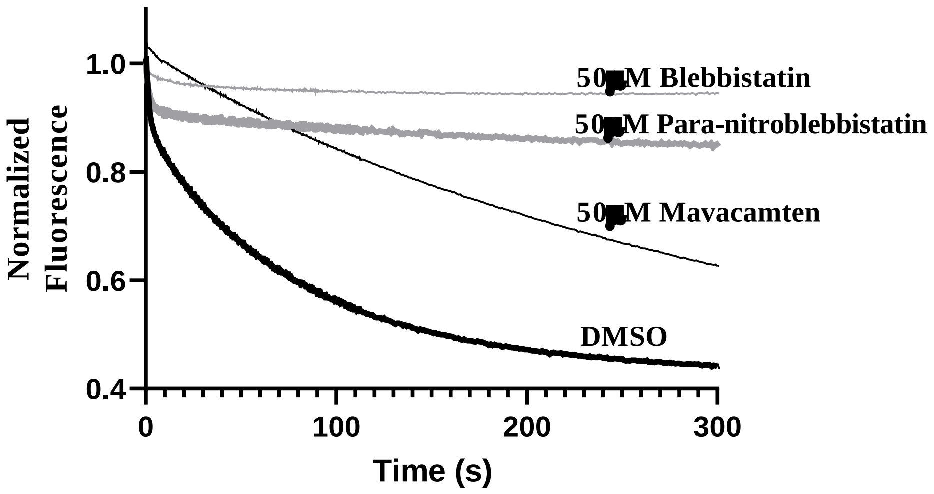


**Figure S4.** Comparing the normalized fluorescence decay profiles measured in bovine cardiac STFs following treatment with high concentration (50 µM) of blebbistatin, para-nitroblebbistatin, and mavacamten.
